# Illegal trade of morphologically distinct populations prior to taxonomic assessment and elevation, with recommendations for future prevention

**DOI:** 10.1101/2020.05.26.116426

**Authors:** Matthijs P. van den Burg, Bruce J. Weissgold

## Abstract

The negative impacts of international wildlife trafficking are well known, and such negative impacts can be even more pronounced for insular species. This dynamic market needs close monitoring, and when novel species appear in the commercial trade relevant authorities should be able to react in order to reduce negative impacts on wild populations. Here we describe a novel case where an insular endemic form of the *Iguana iguana* complex has entered the international commercial trade, likely stimulated by efforts to elevate the form taxonomically. Despite the absence of authorized export permits from the relevant CITES authority, we identify animals that are sold in a range of countries and the likely pathway and methods of importation. We provide recommendations to prevent future illegal collection and trafficking that could be implemented for other taxa. We call for increased awareness of the higher economic value of taxa considered for future taxonomic elevation, and increased monitoring of the commercial trade in order to act promptly when illegal activity is detected.

## Main text

In the current digital era, ongoing efforts to understand and describe Earth’s biodiversity are greatly aided by the increased public availability of scientific information. These efforts are of great interest to researchers and students, policy makers, conservationists, and the general public. However, the entry of certain information into the public sphere can have negative consequences (Stuart, Rhodin, Grismer, & Hansel, 2006; Meijaard, & Nijman, 2014; Lindenmayer, & Scheele, 2017). A major concern is that species descriptions or taxonomic elevations cause wildlife trafficking through increased desirability within the international trade.

One major threat to a variety of species is the global trade in wildlife. Insufficiently regulated and controlled collection and trade can result in unsustainable harvest and subsequent population declines; this is also true for herpetofauna species (Roa, Duckworth, Roberts, & Shepherd, 2014; Auliya, Ariano-Sanchez, Baard, & Brown, 2016; Rowley, Shepherd, Stuart, Nguyen, & Hoang, 2016). As scientists seek to better understand our planet’s biodiversity, novel data can also be used for financial gain through illegal trade, including financial investment speculation in live animals, a newly emergent threat in the commercial reptile trade (Brian Horne, pers. comm). The occurrence of recently (re)discovered reptile species within the pet trade has occurred multiple times (see Auliya et al. [2016] and Rowley et al. [2016]), even when researchers refrained from publishing locality data. For example, the recently described *Echinotriton maxiquadratus* (Mountain spiny crocodile newt) appeared in the pet trade despite the fact that the authors excluded locality data from their original publication and asked others to refrain from sharing the data (Rowley et al., 2016; IUCN & TRAFFIC, 2019). However, specifically stating that locations should not be shared might unintentionally spark interest in the lucrative international trade. Furthermore, it is not unusual for commercial traders to attempt to legalize otherwise illegally collected wild specimens by fraudulently obtaining CITES permits indicating that the animals were bred in captivity (Nijman, & Shepherd, 2009; Janssen, & Chng, 2018). Commercial hobbyists and dealers may be issued CITES (Convention on International Trade in Endangered Species of Wild Fauna and Flora) export permits for the offspring of illegally collected and/or trade animals, therefore, creating a perverse incentive for trafficking. This problem was recently discussed and debated at the 18^th^ meeting of the CITES Conference of the Parties, resulting in an important interpretive statement on the implementation of CITES - Resolution Conf. 18.7, Legal acquisition findings (https://cites.org/sites/default/files/document/E-Res-18-07.pdf). Lastly, although commercial trade in recently described species is common, instances where a proposed taxonomic revision spurs a surge in illegal collection and international trade are rare. This paper summarizes the presence in international commercial trade of specimens of a phenotypic distinct insular population of *Iguana iguana* prior to taxonomic reassessment.

Species of Iguaninae (ITWG, 2016) are long-lived, generally large, herbivorous lizards that occur in the Americas, West Indies and several pacific islands. Globally, Iguaninae are considered among the most endangered lizard groups as >80% of species are considered as threatened (IUCN, 2020), and many species have limited ranges, being insular endemics to one or a small number of islands (ITWG, 2016). Correspondingly, numerous species are listed on Appendix I and II of CITES. As with many other taxa (Hedges et al., 1992), iguanids show a long evolutionary history (Malone, Wheeler, Taylor, & Davis, 2000; ITWG, 2016) and are under current pressure, threatened by a wide range of problems; e.g. invasive species, habitat destruction, hybridization and the impacts of tourism and other anthropogenic activities (Iverson, Grant, Knapp, & Pasachnik, 2016; Pasachnik, Carreras De Leon, & Léon, 2016; van den Burg, Madden, van Wagensveld, & Buma, 2018a; van den Burg, Brisbane, & Knapp, 2020).

*Iguana iguana*, listed as a CITES Appendix II species since 1977 and commonly traded pet (Fig. 1), occurs throughout most of Central- and South America, including a number of Caribbean islands (Bock, Malone, Knapp, Aparicio, & Avila-Pires, 2019). Despite this species’ wide native distribution and identified phenotypic and genetic differences (Lazell, 1973; Malone, & Davis, 2004), extensive phylogeographic and phenotypic diversity research only recently commenced (Breuil, 2013; Stephen, Reynoso, Collett, & Hasbun, 2013; Breuil et al., 2019). However, a thorough study on range-wide phenotypic variation is still lacking. Besides native populations, numerous established alien populations are known, having originated primarily in the pet-trade (Falcón, Ackerman, & Daehler, 2012; van den Burg, van Belleghem, & De Jesús Villanueva, 2020). In fact, the pet trade has been identified as a threat to local mainland populations due to the unsustainable harvest of wild animals (Stephen, Pasachnik, Reuter, Mosig & Ruyle, 2011), and despite an allowable and robust legal trade under CITES, trafficking of the species has been recorded from several countries in Central America Caribbean islands (Stephen et al., 2011; Noseworthy, 2017; Breuil et al., 2019).

**Figure 1.**
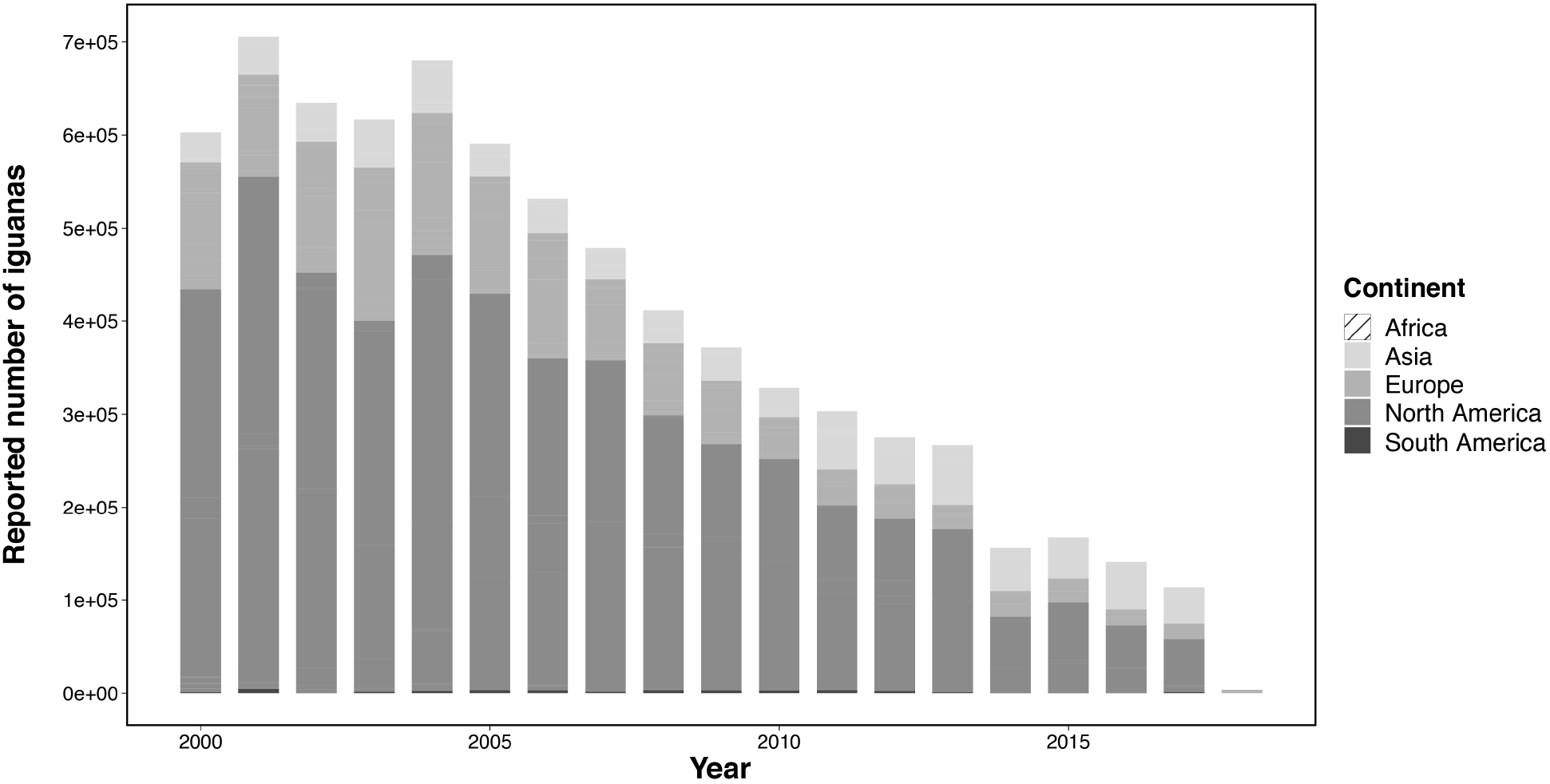
Minimum CITES reported import of Iguana iguana for commercial or personal use 325 between 2000 and 2017.

Based on a morphological assessment of several island localities and mainland *I*. *iguana* lineages, Breuil (2013) suggested the presence of cryptic diversity and the presence of new taxonomic units for the populations on St. Lucia, Saba and Montserrat. Hereafter, Breuil, Vuillaume, Schikorski, Krauss, & Morton (2019) furthered this view: “Breuil et al. (in preparation) work on the genetic and morphological originality of the insular population of Saba and Montserrat”. Additionally, information regarding the intention to taxonomically elevate the Saba population comes from a 2011 social-media post from the Saba Conservation Organization that mentioned: “…a study is [under]way to scientifically establish the Saba Black Iguana as a distinct subspecies of the Green Iguana (*Iguana iguana*), only found on Saba”. This information was shared widely and can be found on numerous websites and social media groups. Overall, information concerning the potential uniqueness and future scientific recognition of the melanistic *I*. *iguana* populations of Saba, and Montserrat, has been publicly available for seven years; and was recently elevated to species level, *Iguana melanoderma* (Breuil, et al. 2020). Here we focus on the trade of these melanistic *Iguana* localities (Saba and Montserrat) and identify countries were such animals are sold, and the pathway and methods of importation and trade; data collation and manuscript preparation were completed before publication of Breuil et al. (2020). We collated information from the CITES Trade Database [https://trade.cites.org/], through on-line reviews of advertisements and social-media posts about melanistic iguanas and contact with commercial hobbyists, dealers and private keepers.

No *I*. *iguana* exports from Saba are reported in the CITES Trade Database through the end of 2019 (CITES, 2020). However, based on data collected in 2016 (Noseworthy, 2017) and our review of online advertisements and social-media posts, animals advertised as “Black dragons,” “Saba black iguanas,” “Saba Island iguanas” with phenotypic characteristics of the melanistic populations are currently in private collections and/or sold in the United States, Germany, Indonesia, Belgium, Malaysia and Japan. We found that at least 12 companies/collectors have acquired iguanas indicated as being from Saba since 2015, with recorded posts and adverts for every year. Furthermore, although to collection of detailed information on wildlife trade routes and business practices can be difficult, we gained insights about the origin and trade pathways from retailers and private owners of melanistic iguanas. Namely, one U.S.A. retailer states online about their melanistic Saba iguanas: “They are captive bred on a different island and imported”; including one image with info keywords “SXM” (Sint Maarten) and “Anguilla”. In addition, one commercial buyer described how melanistic animals from St. Maarten have been imported into Europe as captive-bred animals under CITES permits approved by the local authority. Lastly, another commercial buyer indicated that Saba iguanas are imported into the United States from Barbabos. Overall, despite the absence of CITES export records from Saba (CITES, 2020), all sellers claim to have appropriate CITES documentation accompanied with their stock animals.

Claims of captive breeding have been fraudulently used to trade illegally caught wild animals (Jansen, & Chng, 2018). We tracked official documentation (CITES, 2020) after receiving information about pathways and origins of melanistic iguanas in the trade (see above). We found that captive-bred *I*. *iguana* have been exported from Sint Maarten and Barbados (CITES, 2020), islands without unique species localities but instead alien *I*. *iguana* populations (Censky, Hodge, & Dudley, 1998; van den Burg, Meirmans, van Wagensveld, Kluskens, & Madden, 2018b). Namely, since 2014, a total of 807 live iguanas have been exported from Sint Maarten (n=381) and Barbados (n=426), to Canada, Czech Republic, Germany, Hong Kong, Japan, United States and Thailand. Information requested from these CITES registered permits in June 2019 had not been provided by both the Sint Maarten and Barbados authorities at the time of acceptance of this manuscript, despite previous contact between the authors and the St. Maarten authority. As trade in the CITES Trade Database is reported as *I*. *iguana,* we cannot be certain that all these captive-bred exports are of Saba iguanas, or of the local invasive populations. However, we only found one retailer selling St. Maarten iguanas, and none for Barbados iguanas. Given multiple lines of evidence we argue that illegal trade in iguanas from Saba is facilitated through either false claims of captive breeding, and/or successful breeding of trafficked founder stock on Sint Maarten and Barbados; close regional proximity of St. Maarten and Barbados to Saba and Montserrat, the absence of native *I*. *iguana* populations on St. Maarten and Barbados, only a single retailer appears to sell iguanas from St. Maarten, and based on information about origin and transport pathway from purchasers of melanistic iguanas. We argue that this illegal trade is facilitated through the issuance of CITES export permits by the relevant authorities.

Here we report the illegal international trade of a species locality prior to taxonomic assessment and elevation for the high-end commercial hobbyist pet market. As no exports have been reported by the relevant CITES authorities, we provide recommendations to combat trafficking and better ensure that CITES permits are not issued for the onward trade of smuggled animals and/or their offspring. We recommend that Management Authorities of distinct insular *Iguana* lineages (Saba, Montserrat, St. Lucia, St. Vincent and Grenadines) publish zero annual export quotas to communicate that the commercial export of their populations are not permitted. To ensure that governance or corruption issues are not impacting the regulation and enforcement of applicable laws and regulations, the CITES Secretariat should work with the relevant authorities of St. Maarten to determine why their 2015-2018 *I. iguana* exports were not reported. This will better ensure all trade is legal and properly documented. With ongoing efforts to understand *Iguana* phylogeography, additional divergent phenotypes will likely be discovered. Hence, we recommend that the CITES Animals Committee consider a revision of the *I*. *iguana* Standard Taxonomic Reference to ensure that it reflects currently accepted taxonomy. For CITES-listed species, relevant Management Authorities should only permit export if they are satisfied the specimens and their founder stock were not obtained illegally; the guidance of CITES Resolution Conf. 18.7, <u>Legal acquisition findings</u> can be instructive in these efforts. Especially in phenotypic diverse taxa an identification guide could help efforts to counter smuggling (Stephen, & Binns, 2011) and help exporting and importing Management Authorities to prevent fraudulent trade practices. Additionally, a genetic reference database for variable loci can aid the identification of the origin of ceased individuals, although we note the current logistical, cost and related difficulties in applying such forensic techniques to combat wildlife trafficking. With shifting taxonomy and descriptions of new species, these recommendations are not isolated to the present case and can help prevent illegal trade in current and novel species, including unique localities.

In the situation presented here, as well as in others where the fraudulent use of CITES permitting system is suspected, both exporting and importing countries can take action. Namely, exporting Parties (St. Maarten and Barbados) should review their export permit issuance processes to ensure that permits are not issued for trafficked animals or their offspring. Additionally, importing Parties should pay close attention to imports of *Iguana* sp. of these and other Caribbean islands to better ensure that illegal trade is detected. Importing Parties should scrutinize export permits and contact the relevant authorities (Saba, St. Maarten and Barbados) to verify the authenticity of permits and to ensure that those were issued in compliance with actual CITES standards. When fraudulent use of the CITES permitting system is detected or suspected, the relevant authority (here Saba) should make other CITES Parties aware to better detect irregularities and take actions. One method is to request the CITES Secretariat to issue a Notification to the Parties indicating that no lawful export (of *Iguana melanoderma*) has been authorized, and therefore no Management Authorities should issue (re-) export permits or permits for offspring from such illicit founder stock. This step will strengthen the implementation of CITES Resolution Conf. 18.7, Legal acquisition findings; an important interpretive statement designed to combat this very type of illegal international wildlife trade. Above methods can help prevent negative conservation impacts of illegal wildlife trade and safeguard the integrity of the CITES permitting system, which is intended to prevent the unsustainable exploitation of wild species. Finally, we urge zoos and commercial hobbyists to carefully examine the origins of specimens and their founder stock - before adding new animals to their collections - to better ensure that they were not illegally acquired.

The majority of the ~7.8 million animal species yet-to-be described on Earth will likely be small ranged and rare (Mora, Tittensor, Adl, Simpson, & Worm, 2011), and potentially of great interest to the commercial hobbyist and pet trade, especially those with unique phenotypic characteristics. Researchers should strengthen their awareness of the potential negative consequences associated with publication of sensitive information not only for new or rediscovered taxa but also for those under consideration of taxonomic elevation. Expert advice in cases where future illegal activity can be expected can be sought from the wider wildlife trade community. Prior to publication of sensitive information, researchers should ensure the relevant CITES and enforcement authorities are informed so they are able to monitor for trafficking and can take relevant steps to prevent negative conservation impacts. Globally, it will be important to closely monitor the commercial hobbyist and pet trade and rapidly react to newly traded species in order to prevent overexploitation, extirpation or even extinction.

## Acknowledgements

We thank Catherine Malone and Willow Outhwaite for improving an older version of this manuscript. This research received no specific grant from any funding agency, or commercial or not-for-profit sectors.

